# Base editing in bovine embryos reveals a species-specific role of SOX2 in regulation of pluripotency

**DOI:** 10.1101/2021.11.10.468023

**Authors:** Lei Luo, Yan Shi, Huanan Wang, Zizengchen Wang, Yanna Dang, Shuang Li, Shaohua Wang, Kun Zhang

## Abstract

The emergence of the first three lineages during development are orchestrated by a network of transcription factors, which are best characterized in mice. However, the role and regulation of these factors are not completely conserved in other mammals, including human and cattle. Here, we establish a gene inactivation system by introducing premature codon with cytosine base editor in bovine embryos with a robust efficiency. Of interest, SOX2 is universally localized in early blastocysts but gradually restricted into the inner cell mass in cattle. SOX2 knockout results in a failure of the establishment of pluripotency. Indeed, OCT4 level is significantly reduced and NANOG was barely detectable. Furthermore, the formation of primitive endoderm is compromised with few SOX17 positive cells. Single embryo RNA-seq reveals a dysregulation of 2074 genes, among which 90% are up-regulated in SOX2-null blastocysts. Intriguingly, more than a dozen lineage-specific genes, including *OCT4* and *NANOG*, are down-regulated. Moreover, SOX2 expression is sustained in the trophectoderm in absence of CDX2 in bovine late blastocysts. Overall, we propose that SOX2 is dispensable for OCT4 and NANOG expression and disappearance of SOX2 in the trophectoderm depends on CDX2 in cattle, which are all in sharp contrast with results in mice.

**Significance:** The first and second cell fate decisions of a new life are important for subsequent embryonic and plancental development. These events are finely controlled by a network of transcriptional factors, which are extensively characterized in mice. Species-specific roles of these proteins are emerging in mammals. Here, we develop a gene loss-of-function system by using cytosine base editors in bovine embryos. We find that expression pattern, functional roles, and regulation of SOX2 are all different between mouse and bovine embryos. Remarkbly, SOX2 is required for OCT4 and NANOG, two well established pluripoteny genes. Furthermore, CDX2 is required to shut down SOX2 in the trophectoderm. Given similar expression pattern of SOX2 between human and bovine blastocysts, bovine embryos represents a putative model to investigate human pluripotency regulation in vivo.

## Introduction

Mammalian preimplantation development is characterized of the two earliest cell fate decisions. The first cell fate decision gives rise to the inner cell mass (ICM) and the trophectoderm (TE) and subsequently the ICM generates the primitive endoderm (PE) and the epiblast (EPI) during the second cell fate decision. TE and PE will develop into placenta and extra-embryonic cells, respectively, whereas the pluripotent EPI contributes to the embryo proper (1, 2). The mechansims that regulate these events have been mostly obtained from mouse model. Recent gene-expression and functional analyses suggest that these mechanisms in the mouse may differ in other mammals, including human and cattle. Investigation of these mechansims is important for assisted reproductive technology, regenerative medicine as well as understanding early embryonic mortality in human and agricultural animals.

The establishment and maintenance of pluripotency is regulated by a variety of transcription factors, including core pluripotency factors, OCT4, SOX2 and NANOG (1, 2). The functional importance and relationship of these core transcription factors have been relatively well-characterized in mouse embryos. Interestingly, unlike the universal expression pattern of OCT4 in morula, SOX2 is specifically restricted into the inside cells of morula that become the ICM and is considered the earliest pluripotency marker in mice (3). The HIPPO pathway plays a critical role in temporal and spatial expression of SOX2 in mouse preimplantation embryos (3, 4). However, SOX2 is not required for the first cell fate decision although SOX2-null mouse embryos die soon after implantation and exhibit abnormal ICM (5). Meanwhile, SOX2 is dispensable for intial expression of OCT4 and NANOG in mouse blastocysts (3). However, unlike the expression pattern in mouse preimplantation embryos, SOX2 is not restricted until the expanded blastocyst stage in cattle and humans (6), suggesting a differential regulation of pluripotency in these species.

Base editors are derived from CRISPR/Cas9 genome-editing system and used for precise base editing without DNA double-strand breaks and homology-directed repair (7). Adenine base editors (ABEs) are used to convert an A:T base pair to a G:C base pair (8). Cytosine base editors are used to convert C:G to T:A (9). In addition, a specific function of cytosine base editor is to install a premature stop codon by converting the four codons CAA, CAG, CGA and TGG into stop codons TAA, TAG or TGA (10, 11). To date, base editing has been successfully implemented in the embryos of mice (12, 13), rat (14), pig (15), rabbit (16), cynomolgus monkey (17), and humans (18–20) but has yet been determined in cattle.

In the present study, we successfully develop a high-efficient base editing system in bovine embryos, representing a poweful tool to interrogate gene functions. We then address the role of SOX2 in bovine early embryonic development. Sox2-null embryos can develop to the blastocyst stage, but the ICM is abnormal. SOX2 deletion results in a significant reduction in OCT4 and NANOG expression and dysregulated expression of over 2000 genes. Impressively, CDX2 inhibits SOX2 expression in the TE of bovine late blastocysts. In summary, SOX2 is required for OCT4 and NANOG expression and CDX2 is necessary for ensuring the restricted expression of SOX2 in the ICM of bovine blastocysts.

## Results

### Base editors enable efficient genome editing in bovine embryos

We first sought to establish base editing system using the cytosine base editor (BE3) and the adenine base editor (ABE7.10). Base editor mRNA and sgRNA targeting *SMAD4* were co-injected into the zygotes. Morula cultured in vitro for 6 days were collected and genotypes identified by Sanger sequencing and targeted deep sequencing (Fig 1A and 1B). As for BE3, the desired mutations of C_6_ and C_7_ to T were found in all embryos examined (Fig 1C). The average efficiency of C_6_ and C_7_ being edited as T was 86.3% and 85.4%, respectively, compared with 5.0% in wildtype (WT) embryos (Fig 1D). Regarding ABE7.10, the target mutation of A_5_ to G was also found in all embryos with an average editing efficiency of 79.4% versus 4.5% in WT embryos (Fig 1E and 1F). In addition, we investigated the off-target effect by targeted next-generation sequencing and found no obvious distal off-target edits at the six predicted off-target sites (Fig S1A and S1B).

**Figure 1.**
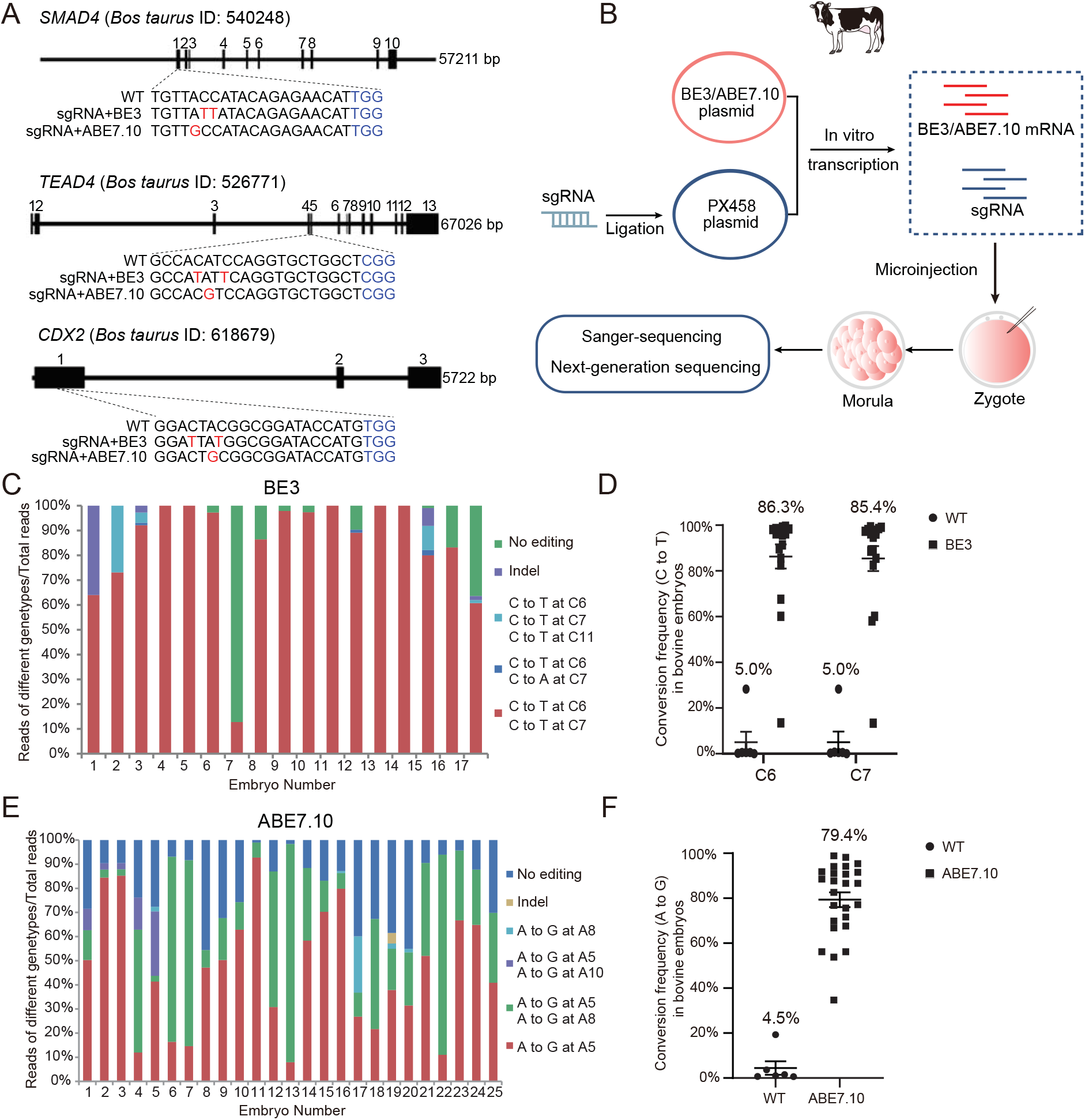
Base editors enable efficient genome editing in bovine embryos. A. Target sites of sgRNA designed for *SMARD4*, *TEAD4*, *CDX2*. Red letters represent the target sites of BE3 or ABE7.10. B. Experimental scheme for base editing in bovine early embryos. C and D: Results of targeted deep sequencing for base editing of SMAD4 by BE3 in bovine embryos. 17 embryos were analyzed. E and F: Results of targeted deep sequencing for base editing of SMAD4 by ABE7.10 in bovine embryos. 25 embryos were analyzed.

To evaluate the ability of base editors to edit multiple genes in bovine embryos, we simultaneously injected sgRNAs targeting three genes, *SMAD4*, *TEAD4*, and *CDX2*, with base editor mRNA. Results showed injection of the base editor components did not affect embryonic development to morula stage (Fig S2A and S2B). Using BE3, results indicated successful editing of three, two and single target genes in 25.8%, 25.8% and 12.9% embryos, respectively (Fig S2C and Table 1). For ABE7.10, results indicated successful editing of three, two and single target genes in 23.3%, 56.7% and 20.0% embryos, respectively (Fig S2D and Table 2). Taken together, these data present proof-of-evidence of base editing with high efficiency in bovine embryos.

**Table 1.** Results of multiple genes’ editing in bovine embryos by BE3

| No. of embryos sequenced | No. of unedited (%) | No. of single gene-edited (%) |  |  | No. of two genes-edited (%) |  |  | No. of three genes-edited (%) |
| --- | --- | --- | --- | --- | --- | --- | --- | --- |
|  |  | SMAD4 | TEAD4 | CDX2 | SMAD4 and TEAD4 | TEAD4 and CDX2 | SMAD4 and CDX2 | SMAD4, TEAD4 and CDX2 |
| 31 | 11 (33.5) | 2 (6.5) | 1 (3.2) | 1 (3.2) | 0 (0.0) | 1 (3.2) | 7 (22.6) | 8 (25.8) |

**Table 2.** Results of multiple genes’ editing in bovine embryos by ABE7.10

| No. embryos sequenced | No. unedited (%) | No. of single gene-edited (%) |  |  | No. of two gene-edited (%) |  |  | No. of three gene-edited (%) |
| --- | --- | --- | --- | --- | --- | --- | --- | --- |
|  |  | SMAD4 | TEAD4 | CDX2 | SMAD4 and TEAD4 | TEAD4 and CDX2 | SMAD4 and CDX2 | SMAD4, TEAD4 and CDX2 |
| 30 | 0 (0.0) | 0 (0.0) | 1 (3.3) | 5 (16.7) | 1 (3.3) | 1 (3.3) | 15 (50.0) | 7 (23.3) |

### Disrupting genes by introducing premature stop codon with cytosine base editor in bovine embryos

Cytosine base editors can introduce premature stop codons to inactivate genes by precisely converting four codons into stop codons. We next test the feasibility of disrupting a gene in bovine early embryos. To maximize the editing efficiency, we designed and simultaneously microinjected 2 sgRNAs targeting the gene of interest, *OCT4* (Fig S3A). Results shows the edited efficiency of sgRNA1 is 25.92% and sgRNA2 reaches 74.07% (Fig S3B and S3C). Overall, premature stop condons were successfully introduced in 77.8% (21 out of 27) embryos, suggesting gene disruption. As a side-by-side experiment, immunostaining analysis confirmed that OCT4 can be efficiently deleted in all blastomeres in these blastocysts (Fig S3D and S3E). Next, we tested if OCT4-null embryos generated here recapitulate the phenotype of OCT4 knockout embryos produced via somatic cell nuclear transfer as reported previously (21). Remarkbaly, NANOG is barely detectable in absence of OCT4 at blastocyst stage (Fig S3D and S3F). The developmental potential to form blastocysts is grealty inhibited in OCT4 KO groups (Fig S3G). These results are consistent with the previous study. Thus, we establish a powerful and reliable system to accomplish efficient base editing in bovine embryos, which will facilitate studies of gene functions in bovine embryos.

### Expression pattern of SOX2 protein in bovine early embryos

To functionally characterize SOX2 in bovine embryos, we first determine its expression pattern in detail in bovine embryos. SOX2 was first found in the 8-cell stage and continued to express thereafter (Fig 2A). In contrast to mouse embryos, SOX2 was not detected during oocyte maturation and the early development to the four-cell stage (Fig 2A). It is noteworthy that SOX2 gradually accumulates in the ICM cells along with blastocyst expansion. Specifically, SOX2 was evenly distributed in both TE and ICM in early blastocysts. Then, SOX2 was lost in subsets of TE cells in middle blastocysts, and eventually restricted into ICM in late blastocysts (Fig 2B). Quantitative results showed SOX2 level in TE is gradually diminished relative to the one in ICM when the blastocyst is expanding (Fig 2C and 2D). Altogether, these data indicate that SOX2 displays a different expression pattern in bovine embryos, which suggests that there are differences in the regulatory mechanism of pluripotency between species.

**Figure 2.**
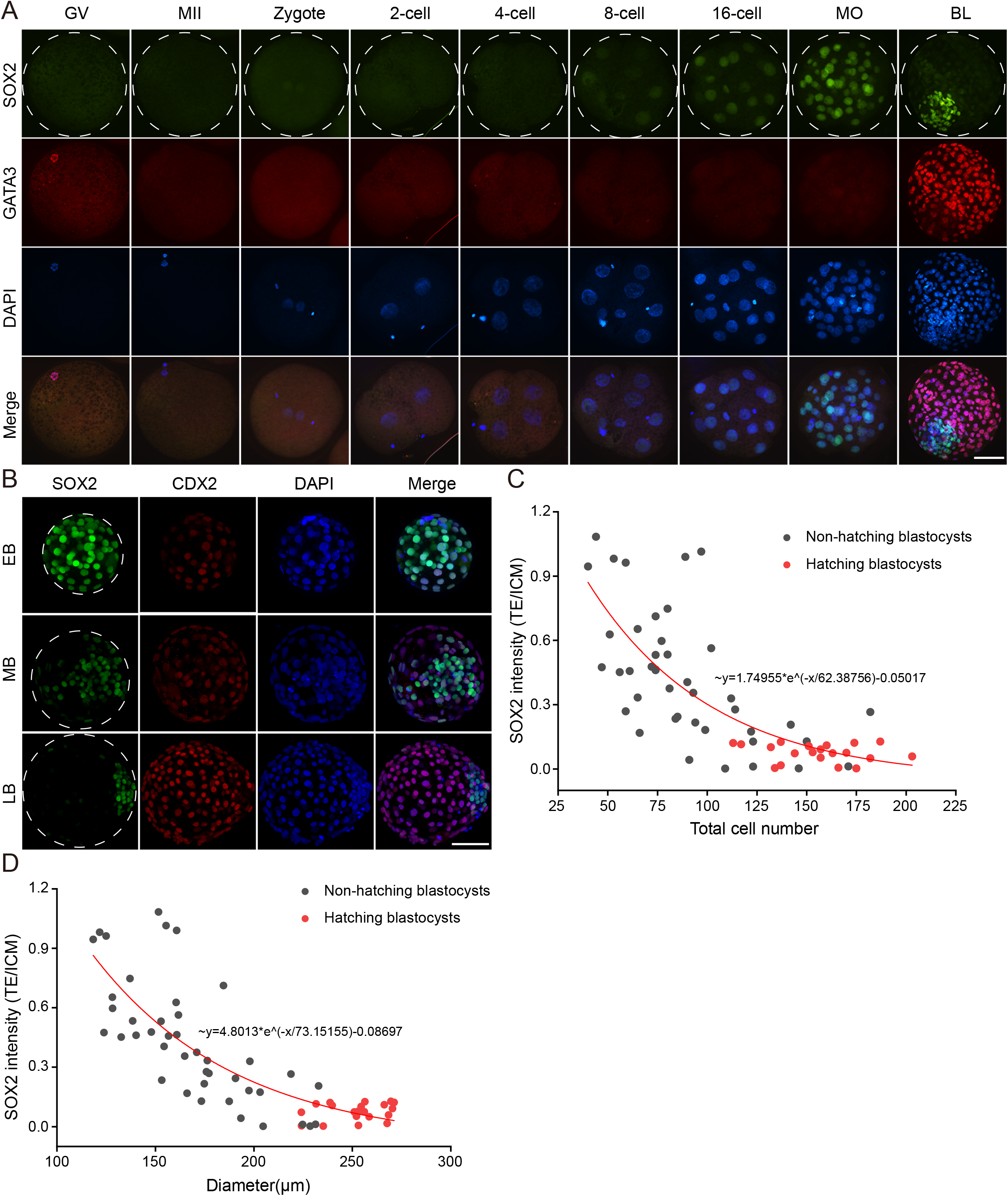
Dynamic expression pattern of SOX2 during bovine early embryonic development. A. Immunofluorescence detection of SOX2 and GATA3 during oocyte maturation and embryonic development. Green: SOX2 protein; Red: GATA3 protein; Blue: DAPI (Nuclei). The experiment was independently replicated two times with at least 10 oocytes or embryos per stage analyzed. Scale bar = 50 μm. GV: germinal vesicle, MII: metaphase II, MO: morula, BL: blastocysts. B. Immunofluorescence analysis of SOX2 protein along with blastocyst expansion. Green: SOX2 protein; Red: CDX2 protein; Scale bar = 50 μm. EB: early blastocyst, MB: middle blastocyst, LB: late blastocyst. C and D: The correlation between SOX2 intensity (TE/ICM) and total cell number or diameter. Note: the red bullets represent hatching blastocysts.

### Effects of SOX2 KO on the bovine embryo development

We then sought to explore the functional role of SOX2 by disrupting its expression using BE3 (Fig 3A and 3B). Genotyping results show sgRNA2 and 3 are more efficient than sgRNA1 in editing SOX2 (Fig 3C and 3D). Overall, premature stop condon was successfully installed at SOX2 in 87.1% (101 out of 116) bovine blastocysts when these three sgRNAs were injected. Immunostaining results further confirmed that SOX2 signal was drastically diminished in these corresponding blastocysts and only 9 (out of 116) embryos display mosacism (Fig 3E).

**Figure 3.**
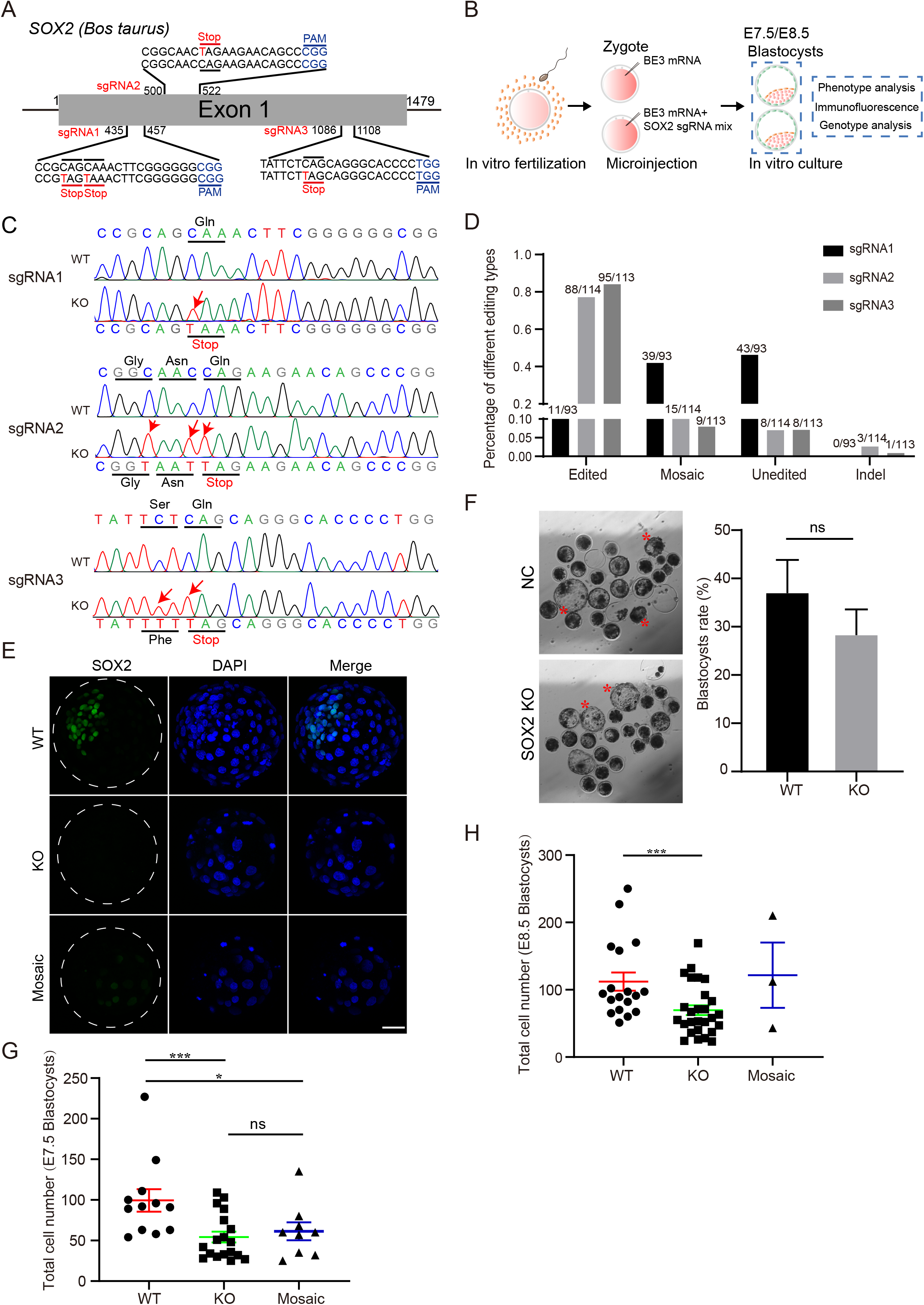
Effects of SOX2 knockout on bovine early embryonic development. A. sgRNAs used to target *SOX2*. Red lines represent the position of introduced premature stop codon. B. Experimental design to explore the effects of SOX2 KO on bovine early embryonic development. C. Representative genotyping results for three distinct sgRNA. WT: wide type; KO: putative SOX2 knockout embryos. Red arrows denote successful C: T conversion. D. Statistical analysis of editing efficiency for each sgRNA. The target sequence of sgRNA1, sgRNA2 and sgRNA3 were analyzed in 93, 113 and 114 embryos, respectively. E. Immunostaining validation for SOX2 knockout at blastocyst (BL) stages (Three replicates of 3-5 embryos were analyzed per group). Scale bar = 50 μm. F. Blastocyst formation rate of bovine embryos after SOX2 knockout. The rate of blastocysts at E7.5 was recorded with no significant difference found between WT and KO groups (Five independent replicates of 20-25 embryos per group). Red asterisks represent hatching blastocysts. Scale bar = 100 μm. G and H: Statistical analysis of total cell numbers at E7.5D (G) and E8.5D (H). Asterisks refer to significant differences (*:P < 0.05; **:P <0.05; ***: P<0.001).

In vitro culture of embryos revealed no significant difference in the capability to become blastocysts between SOX2 KO and WT groups (Fig 3F). Interestingly, the total cell number per blastocyst was significantly reduced at both E7.5 and E8.5 (Fig 3G and 3H). However, the TE cell number (CDX2 positive) was not obviously changed while the number of ICM cells (CDX2 negative) was rather decreased dramatically (Fig 4A). Overall, SOX2 is not required for blastocyst formation, but essential for the ICM development in bovine blastocysts.

**Figure 4.**
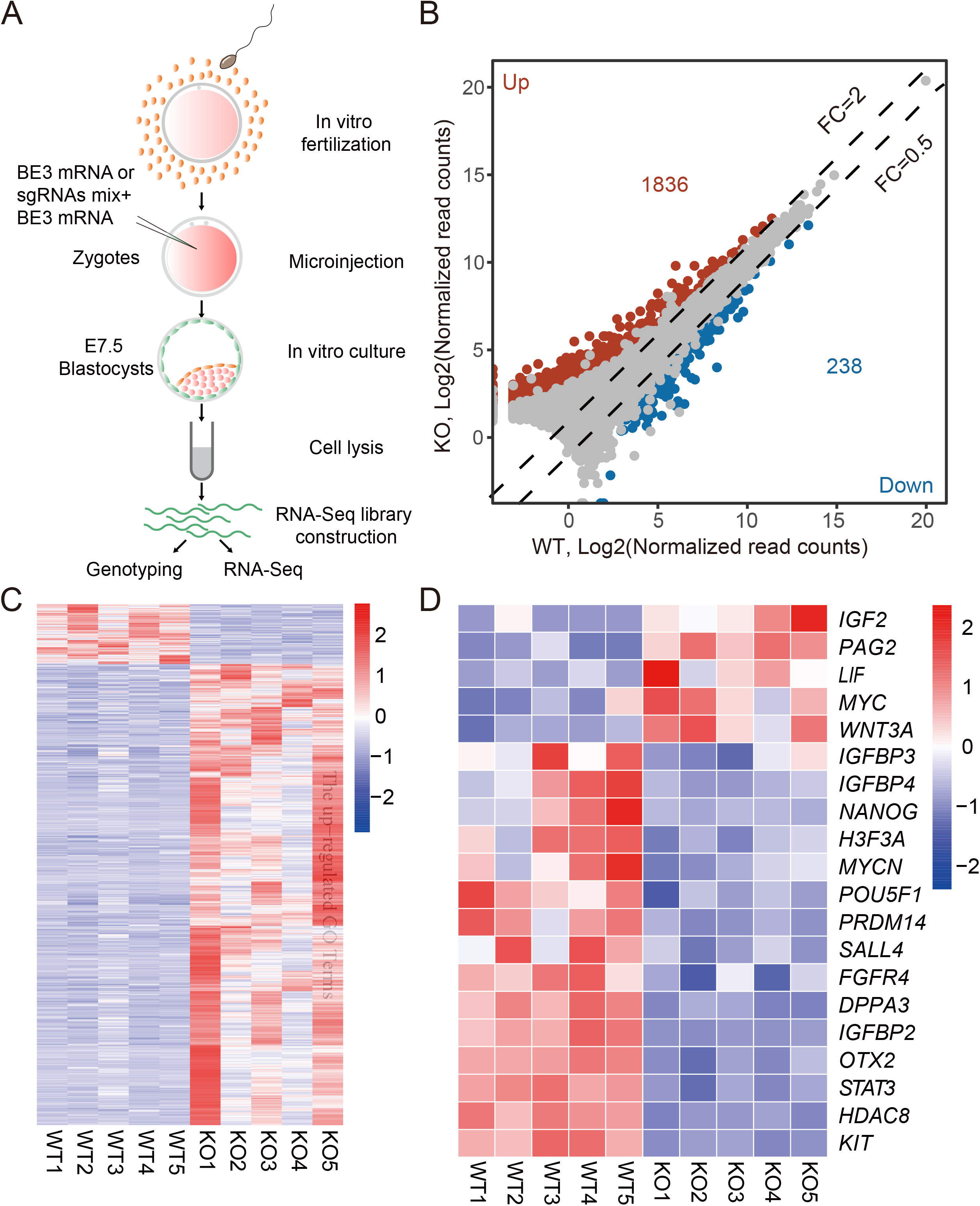
SOX2 knockout disrupts the network of pluripotent genes in bovine blastocysts. A. Experimental scheme of single blastocyst RNA-seq in WT and SOX2 KO groups. B. Volcano plot depicting differentially expressed genes, among which 1836 are upregulated and 238 are downregulated (Fold Change>2 or <0.5; P adj< 0.05). C. Heat map showing all differentially expressed genes between WT and SOX2 KO groups. D. Heat map showing differential expression of genes involved in regulation of pluripotentcy between WT and SOX2 KO group.

### SOX2 knockout disrupts the network of pluripotent genes in bovine blastocysts

To determine the molecular consequence of SOX2 KO, RNA-seq of single blastocyst was performed at E7.5. The samples are morphological indifferent upon collected to avoid bias (Fig 4A and S4A). Partial cDNA libarary from each blastocyst was used to determine the genotype (Fig 4B and S4B). Principal component analysis (PCA) showed that WT and SOX2 KO blastocysts formed two distinguished clusters (Fig S4C). There are a total of 2074 differentially expressed genes (DEGs, Fold changes (FC) >2 or <0.5, P adjusted<0.05), among which 88.53% were remarkably upregulated in SOX2 KO groups (Fig 4B and 4C).

Gene ontology (GO) analysis revealed that the top GO terms enriched in DEGs include membrane depolarization during an action potential, cell adhesion, integral component of plasma membrance, calcium ion binding. Interestingly, we found a number of overrepresented genes that involved in regulation of the pluripotency, including up-regulated *IGF2*, *PAG2*, *LIF*, *MYC*, *WNT3A*, and down-regulated *IGFBP3*, *IGFBP4*, *PRDM14*, *SALL4*, *FGFR4*, *STAT3*, *HDAC8* (Fig 4D). Surprisingly, *NANOG* and *OCT4* were both sharply downregulated in SOX2 KO bovine blastocysts. In sum, these data suggest SOX2 plays a critical role in maintaining correct gene expression of the pluripotency network.

### SOX2 is indispensable for NANOG and OCT4 expression in the ICM of bovine blastocysts

We then hypothesized that SOX2 is required for OCT4 and NANOG expression in bovine blastocysts. As reported previously, we confirmed OCT4 is evenly localized in the ICM and TE at early and middle blastocyst stage but gradually restricted into the ICM at late blastocyst stage in cattle (Fig S5A and S5B). Remarkably, the intensity of OCT4 decreased significantly in the SOX2 KO blastocysts (Fig 5B). NANOG was first detected in the morula stage and distributed in both TE and ICM of the early blastocyst and then fast aggregated into epiblast in the late blastocyst stage (Fig S5B). Intriguingly, NANOG was barely seen in SOX2 KO blastocysts (Fig 5C). These results collectively suggest that SOX2 is the core gene and upstream of OCT4 and NANOG in the network of pluripotency genes in cattle.

**Figure 5.**
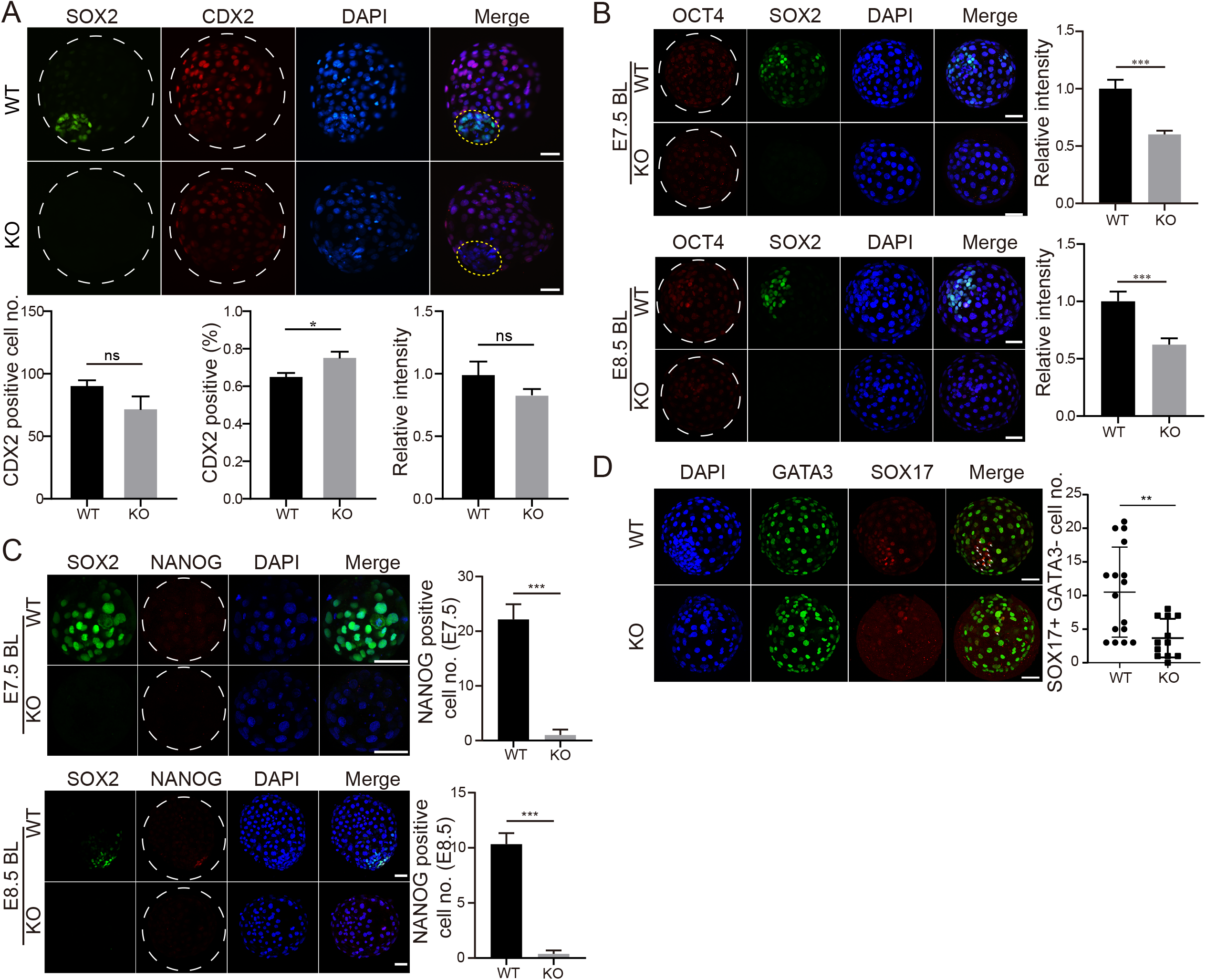
SOX2 is indispensable for NANOG and OCT4 expression in the ICM of bovine blastocysts. A. Immunostaining analysis of CDX2, a marker of trophectoderm (TE), in WT and SOX2 KO blastocysts at E8.5. B. Total cell counting analysis of TE cells (CDX2^+^) in WT and SOX2 KO blastocysts (Three replicates of 6-8 blastocysts per group were analyzed). B. Immunostaining analysis of OCT4, a pluripotency marker, in WT and SOX2 KO blastocysts at E7.5 and E8.5 (Two independent replicates of 12-17 embryos per group were analyzed at E7.5 and E8.5, respectively). C. Immunostaining analysis of NANOG, an epiblast marker, in WT and SOX2 KO blastocysts at E7.5 and E8.5 (Two independent replicates of 13-14 embryos per group were analyzed at E7.5 and E8.5 days, respectively). M: Immunostaining analysis of GATA3 (a marker for trophectoderm) and SOX17 (a marker for primitive endoderm) in WT and SOX2 KO blastocysts (Two independent replicates of 9-10 embryos per group were analyzed at E7.5 and E8.5 days, respectively). Asterisks refer to significant differences (*:P < 0.05; **:P <0.05;***: P<0.001). Scale bar = 50 μm.

To further determine if SOX2 KO affect the specification of primitive endoderm, we performed immunostaining against SOX17 and found the number of SOX17 positive cells was greatly reduced after SOX KO (Fig 5D), suggesting a compromised primitive endoderm.

### CDX2 is required for the restricted expression of SOX2 in the ICM of bovine late blastocysts

We next asked if CDX2 is involved in the gradual disappearance of SOX2 in the TE. Both genotyping and immunostaining results indicated CDX2 is completely knocked out in 82.9% embryos (58 out 70; Fig S6A-D). No difference was found on the developmental potential to arrive blastocyst stage in CDX2 KO groups (Fig S6E). Immunostaining analysis revealed that SOX2 signal sustained in the TE of bovine late blastocysts in CDX2 KO groups (Fig 6A-C). To further test the specificity of the role of CDX2 in regulating SOX2 expression during the bovine embryonic development, we microinjected base editor components into one blastomere at 2-cell stage (Fig 6D). Immunostaining and confocal microscopy analysis indicated that SOX2 signal of CDX2 negative cells are obviously brighter than those of CDX2 positive cells in the TE of bovine blastocysts (Fig 6E and 6F), further consolidating the conclusion that CDX2 is required to diminish SOX2 in the TE of bovine late blasotcysts.

**Figure 6.**
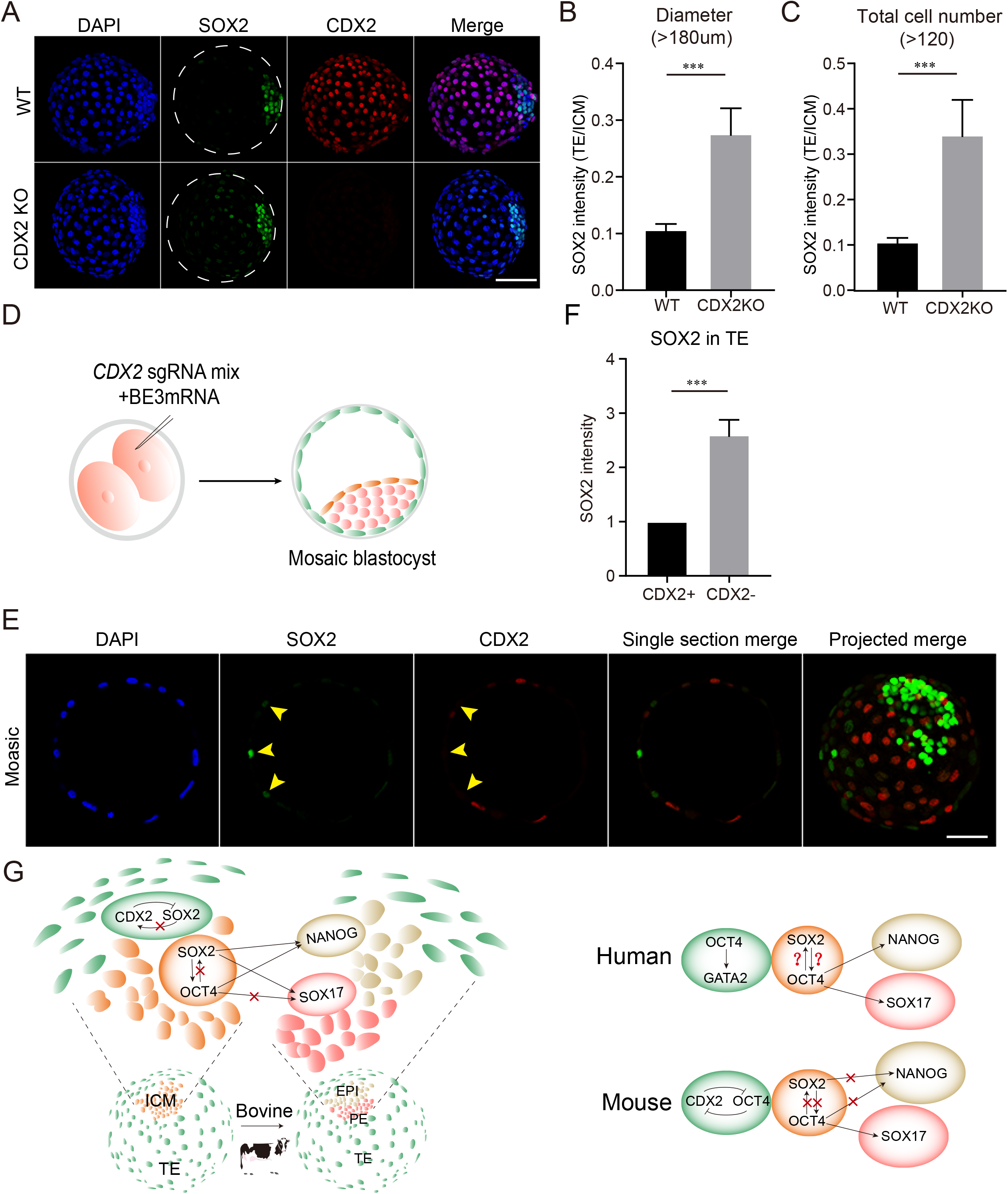
CDX2 is required for the restricted expression of SOX2 in the ICM of bovine late blastocysts. A. Immunostaining analysis of SOX2 in trophectoderm between WT and CDX2 KO blastocysts. Scale bar = 50 μm. Seven replicates of 4-8 blastocysts per group. B and C: Statistical analysis of SOX2 levels (TE/ICM) between WT and CDX2 KO groups when blastocysts diameter >180 μm (B) and total cell number >120 (C). n=23. Asterisks refer to significant differences (***: P<0.001). D. Experimental scheme to produce CDX2 mosaic bovine blastocysts. E. Immunostaining analysis of SOX2 levels in CDX2^-^ TE cells relative to CDX2^+^ TE cells. Green: SOX2 protein; Red: CDX2 protein; Scale bar = 50 μm. F. Statistical analysis of SOX2 intensity in CDX2^-^ TE cells relative to CDX2^+^ TE cells. n=12. Asterisks refer to significant differences (*:P < 0.05; **:P <0.05; ***: P<0.001). G. Summary and working model of functional relationship between SOX2 and other core lineage-specific genes in human, mouse and bovine blastocysts.

## Discussion

How the earliest cell fate decisions are made is a fundamental question due to their importance for the establishment of pregnancy and fetus development in mammals. Recent studies suggest specie-specific regulation of these events. Here, we present the proof-of-evidence of base-editor-mediated gene knockout with a robust efficiency in bovine early embryos. Using this platform, we find SOX2 regulates OCT4 and NANOG expression and the disappearance of SOX2 in the TE of bovine blastocyst is dependent on CDX2. These results are different with those of mouse studies, revealing a species-specific role and regulation of SOX2 in mammals.

A longstanding barrier for studying gene functions in large animal is the lack of genetic tool to disrupt a gene of interest. The advent of CRISPR-Cas9 technology represents a powerful approach to achieve genome editing. However, recent studies indicate the use of Cas9 in human early embryos results in unintentional deletion of large fragments, raising concerns in addressing gene functional studies and clinical use (22, 23). We thus decided to use base editing system in the present study. Cytosine base editors are particularly useful for disrupting genes by introducing a premature stop codon into a gene of interest without creating double strand breaks or indels.

Here, our studies reported that BE3 and ABE7.10 enable us to achieve gene editing with an efficiency above 79% in bovine embryos. Importantly, we found no obvious off-target editing at potential site, indicating the specific effects we documented in the present study. To maximize the editing efficieny, we microinjected 2 or 3 sgRNA together and found the target gene can be deleted completely in all blastomeres in around 80% embryos with only less than 10% embryos exhibit mosaicism. We believe this approach is a powerful tool to dissect gene function and produce genome-edited cattle.

A series of transcription factors participate in the lineage development, including CDX2, SOX2, OCT4, and NANOG. These factors are originally identified as lineage-specific in mice and are also present in other mammals. Nonetheless, if their function is conserved across species remains poorly determined, especially in large animals. OCT4 expression lasts a long time in the TE of bovine blastocyst, in contrast to the expression pattern observed in mice (24, 25). OCT4 was restricted into the ICM later than SOX2, suggesting SOX2 is required first for the establishment of pluripotency. Consistenly, OCT4 KO does not affect the expression of SOX2, however, SOX2 KO leads to reduced OCT4 expression in bovine blastocysts. More remarkably, we observed even no NANOG expression in both early or late SOX2-null blastocysts. A recent study show OCT4 is required for NANOG expression in bovine blastocysts (21), suggesting that SOX2 may regulate NANOG indirectly through OCT4. However, we speculate SOX2 also directly regulate the expression of NANOG because OCT4 is not completely lost when SOX2 is deleted (Fig 6G).

RNA-seq results revealed a large-scale disruption of the transcriptome upon SOX2 deletion in bovine blastocysts with 2074 genes expression affected. In comparison, a previous report have shown that only 472 genes are dysregulated in OCT4-null bovine blastocysts (21), indicating the molecular consequence of SOX2 KO is more severe than OCT4.

Mutual feedback between lineage-specific genes in mammalian embryos has been reported previously (1). OCT4 and CDX2 are mutually regulated by each other in the ICM and TE in mouse blastocysts (1). However, Sox2 is restricted to inside cells by a Cdx2-independent mechanism (3). Results herein show CDX2 in involved in suppressing SOX2 in the TE along with blastocyst expansion. This suppression is developmental context-dependent as it takes places in late blastocysts but not in early blastocyst stage.

In conclusion, we demonstrated that the base editing system could be applied to bovine embryos. With this powerful tool, SOX2 knockout was successfully achieved in bovine embryos. Functional experiments proved that SOX2 knockout significantly disrupt OCT4 and NANOG expression. Meanwhile, the disappearance of SOX2 in the TE is dependent on CDX2. Altogether, our study reveals a species-specific role of SOX2 in regulation of pluripotency and unique regulation of SOX2’s restricted expression in blastocysts.

## Materials and Methods

### Materials

All chemicals and reagents were commercially obtained from Sigma (St. Louis, MO) unless stated elsewhere.

### In vitro production of bovine embryos

Bovine embryo in vitro production, including in vitro maturation (IVM), in vitro fertilization (IVF) and in vitro culture (IVC) was performed as procedures published previously with slight modifications (26–28). Briefly, cumulus-oocyte complexes (COCs) containing intact cumulus cells were collected from bovine ovaries obtained from a local abattoir. COCs were matured in Medium-199 (M4530) supplemented with 10% FBS (Gibco-BRL, Grand Island, NY), 1 IU/ml FSH (Sansheng Biological Technology, Ningbo, China), 0.1 IU/ml LH (Solarbio, Beijing, China), 1 mM Sodium Pyruvate (Thermo Fisher Scientific, Waltham, MA), 2.5 mM GlutaMAX™ (Thermo Fisher Scientific, Waltham, MA), and 10 μg/mL Gentamicin at 38.5°C under 5% CO_2_ in humidified air for 22-24 hrs. COCs (60-100 COCs per well in 4-well plates) were then incubated with spermatozoa (1-5×10^6^) purified from frozen-thawed semen by using a Percoll gradient in BO-IVF medium (IVF bioscience, Falmouth, Cornwall, UK). IVF condition was 38.5°C under 5% CO_2_ for 9-12 hrs. Putative zygotes were then removed of cumulus cells by pipetting up and down using Medium-199 (M7528) supplemented with 2% FBS (Gibco-BRL, Grand Island, NY). Embryos were incubated in BO-IVC medium (IVF bioscience, Falmouth, Cornwall, UK) at 38.5°C under 5% CO_2_ in humidified air until use.

### sgRNA design, synthesis, and plasmid construction

BE-Designer online software (http://www.rgenome.net) was used to design sgRNAs. sgRNA sequences with appropriate GC content and low off-target probability were selected that target the protein-coding region of the gene of interest. The sticky end of BpiI: 5’-3’ CACC and 5’-3’ AAAC were added to the 5’ ends of the sense and antisense strand, respectively (Table S1). The DNA sequences were synthesized by Sangon Co., LTD (Shanghai). Then, sgRNA DNA oligos were annealed and cloned into a PX458 vector containing BpiI restriction sites with T7 promoter.

### In vitro transcription

BE3 and ABE7.10 plasmids were purchased from Addgene (#73021 and #102919). After linearization with Not I, the plasmid underwent in vitro transcription using mMESSEAGE mMACHINE T7 kit (Invitrogen) and were purified by LiCl precipitation. sgRNAs were amplified and transcribed in vitro using MEGAshortscript T7 High Yield Transcription Kit (Invitrogen) according to manufacturer’s instructions. Primers are listed in Table S2. After transcription, sgRNAs were purified by ethanol precipitation.

### Microinjection of base editor mRNA

10-20 pL mixture of 100 ng/μL sgRNA and 200 ng/μL ABE7.10 or BE3 mRNA were microinjected into bovine zygotes at 12 h post insemination (hpi) by using a micromanipulator. Control embryos were injected with same amout of mRNA without sgRNA. To maximize the editing efficiency of the gene of interest, a cocktail of two or three sgRNAs were microinjected together with BE3 mRNA.

### Single bovine embryo PCR and genotyping

Injected embryos were collected at morula or blastocyst stage. Genomic DNA was isolated using an embryo lysis buffer (40nM Tris-HCl, 1% Triton X-100, 1% NP-40 and 0.4 ng/mL Proteinase K) at 55 °C for 1 h and 95 °C for 10 min. Nested PCR was performed and then the amplicon was subject to Sanger sequencing. All primers used are listed in Table S3.

### Targeted deep sequencing

Single embryo was subject to whole-genome amplification by using REPLI-g Mini Kits (QIAGEN, Cat. No. 150023). The target sites and 6 potential off-target sites (Table S4) that predicted by an online software were amplified using PCR primers with barcode sequence (Table S5). All amplicons were purified and subject to targeted deep sequencing.

### Immunofluorescence (IF)

Early embryos were rinsed three times with 0.1% PBS/PVP (polyvinylpyrrolidone), and fixed with 4% paraformaldehyde in PBS for 30 min, permeabilized with 0.5% TritonX-100/PBS for 30 min. Fixed samples were then blocked for 1-2 hrs with the buffer containing 10% FBS and 0.1% TritonX-100/PBS. Samples were incubated with primary antibodies for 2 hrs at room temperature or overnight at 4°C. Then, embryos were treated with secondary antibodies for 2 hrs. Nuclear DNA was counterstained by DAPI for 15 min. Samples were mounted and observed with either an inverted epifluorescence microscope (Nikon, Chiyoda, Japan) or a Zeiss LSM880 confocal microscope system (Zeiss, Oberkochen, Germany). For confocal microscopy, Z-stacks were imaged with 5 μm intervals between optical sections. Stacks were projected by maximum intensity to display signals of all blastomeres in one image. All antibody information was shown in Table S6.

### Single blastocyst RNA-seq and data analysis

Single blastocyst from WT and KO group was collected on E7.5. The zona pellucidae of blastocysts was discarded with 0.5% pronase E. The RNA-seq libraries were constructed according to Smart-seq2 procedure as previously described (29). In brief, polyadenylated RNAs were captured and reverse transcribed with Oligo(dT) primer, then the cDNA was pre-amplified using KAPA HiFi HotStart ReadyMix (kk2601). Pre-amplified cDNA was purified with Ampure XP beads (1:1 ratio) and fragmented by Tn5 enzyme (Vazyme, TD502). PCR amplification for 15-18 cycles was performed to prepare sequencing libraries, which were subject to paired-end 150 bp sequencing on a NovaSeq (Illumina) platform by Novogene. The raw sequencing reads were trimmed with Trimmomatic (version 0.39) (30) to generate clean data, and mapped to ARS-UCD1.2 with Hisat2 (version 2.1.0) (31). The raw counts were calculated with featureCounts (version 1.6.3) (32) and underwent differential expression analysis using DESeq2 (33). The differentially-expressed genes between WT and KO group were identified when Padj <=0.05 and Foldchange >=2 or <= 0.5. FPKM for each sample was calculated with Cufflinks (34) for heatmap visualization, and heatmaps generated using pheatmap package in R. Gene ontology analysis was performed with the Database for Annotation, Visualization and Integrated Discovery (DAVID)(35, 36)

### Statistical Analysis

All experiments were replicated at least three times unless stated. Two-tailed unpaired student t-tests were used to compare differences between two groups. The fluorescent intensity was analyzed using Image J as described previously (26). Briefly, the nuclear region was encircled based on the DAPI signal and the intensity measured. The same region was moved to the cytoplasm area and background intensity obtained. The specific signal was calculated by subtracting the cytoplasmic intensity from the nuclear intensity. Finally, the data were normalized to the relative channels in control groups. The graphs were constructed by GraphPad Prism 8.0 (GraphPad Software, USA). P<0.05 refers statistical significance.

## Acknowledgments

We thank all members of the K. Zhang laboratories for their helpful discussions. This work was supported by National Natural Science Foundation of China (No. 31872348, No. 31672416, and No. 32072731 to K.Z.; No. 32072939 to H.W.; No.31941007 to L.L. and S.W.), Zhejiang Provincial Natural Science Foundation (LZ21C170001 to K.Z.; No. LY19C180002 to H.W.) and China Postdoctoral Science Foundation (No. 2020M671742 to L.L.).

---

**Figure S1. Analysis of off-target effects of base editor ABE7.10 and BE3.**

A and B: Targeted deep sequencing analysis of 6 potential off-target sites for ABE7.10 (A) and BE3 (B).

**Figure S2. Application of multi-gene base editing using ABE7.10 and BE3**

A and B: Embryonic developmental rate to reach morula stage in the WT and ABE7.10 (A) or BE3 (B) group (Two replicates of 13-18 embryos per group). C. Representative Sanger sequencing results of ABE7.10-meidated base editing. D. Representative Sanger sequencing results of BE3-mediated base editing. The red letters and frames represent the edited sites. The green letters represent PAM sequence. S-gRNA: *SMAD4* sgRNA; T-gRNA: *TEAD4* sgRNA; C-gRNA: *CDX2* sgRNA.

**Figure S3. Effects of OCT4 knockout on bovine early embryonic development.**

A. Two sgRNAs designed to target *OCT4*. The red letters represent potential editing sites. B. Representative Sanger sequencing results. The red letters represent edited sites. C. Editing types of *OCT4* sgRNA1 and sgRNA2 (27 embryos were analyzed). D. Immunostaining detection of OCT4 and NANOG in WT and OCT4 KO groups (Two replicates of 4-6 blastocysts per group). Green: NANOG; Red: OCT4. Scale bar = 50 μm. E: OCT4 KO results in the decrease of blastocyst rate (Three replicates of 20-25 embryos per group). Scale bar =100 μm. F. Immunostaining analysis of SOX2 expression and distribution (Two replicates of 4-6 blastocysts per group). Green: SOX2; Red: OCT4. Scale bar = 50 μm.

**Figure S4. Single blastocyst RNA sequencing**

A. Single blastocyst was collected (n=5 per group) to perform RNA sequencing. B. Validation of the genotypes of embryos used for RNA-sequencing in A. C. Principal component analysis (PCA) shows high correlation among samples in the same group.

**Figure S5. SOX2 knockout did not affect blastocyst expansion**

A. Quantification analysis of the changes of OCT4 levels in TE and ICM as blastocyst expansion. TE: SOX2^-^ cells; ICM: SOX2^+^ cells. B. The dynamics of NANOG and OCT4 expression accompanied by the blastocyst expansion. Green: NANOG; Red: OCT4 (Two replicates of 3-5 blastocysts at different stages). Scale bar = 50 μm C. SOX2 knockout did not affect the blastocyst expansion at E7.5 or E8.5 (Two independent replicates of 12-17 embryos per group).

**Figure S6. CDX2 KO did not affect bovine early embryonic development.**

A. sgRNAs designed to target CDX2. B. Representative Sanger sequencing results of CDX2 editing. The red letters represent edited sites. C. Immunostaining detection of CDX2 (Three replicates of 5-8 blastocysts per group). Red: CDX2. Scale bar = 50 μm. D. Editing types analysis using *CDX2* sgRNA1, sgRNA2 and sgRNA3 (61 embryos were detected). E and F. CDX2 KO has no effect on the rate of blastocyst formation (Ten replicates of 20-25 embryos per group). Scale bar = 100 μm.

