## Supplemental information for "Base editing in bovine embryos reveals a species-specific role of SOX2 in regulation of pluripotency"

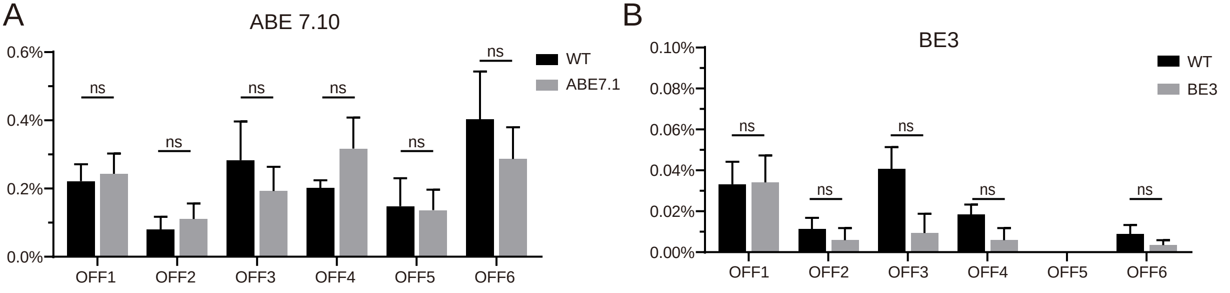
A
ABE 7.10
0.6%
0.4%
0.2%
0.0%
ns
ns
ns
ns
ns
ns
WT
ABE7.1
OFF1
OFF2
OFF3
OFF4
OFF5
OFF6
B
BE3
0.10%
0.08%
0.06%
0.04%
0.02%
0.00%
ns
ns
ns
ns
ns
WT
BE3
OFF1
OFF2
OFF3
OFF4
OFF5
OFF6

**Figure S1. Analysis of off-target effects of base editor ABE7.10 and BE3.**

A and B: Targeted deep sequencing analysis of 6 potential off-target sites for ABE7.10 (A) and BE3 (B).

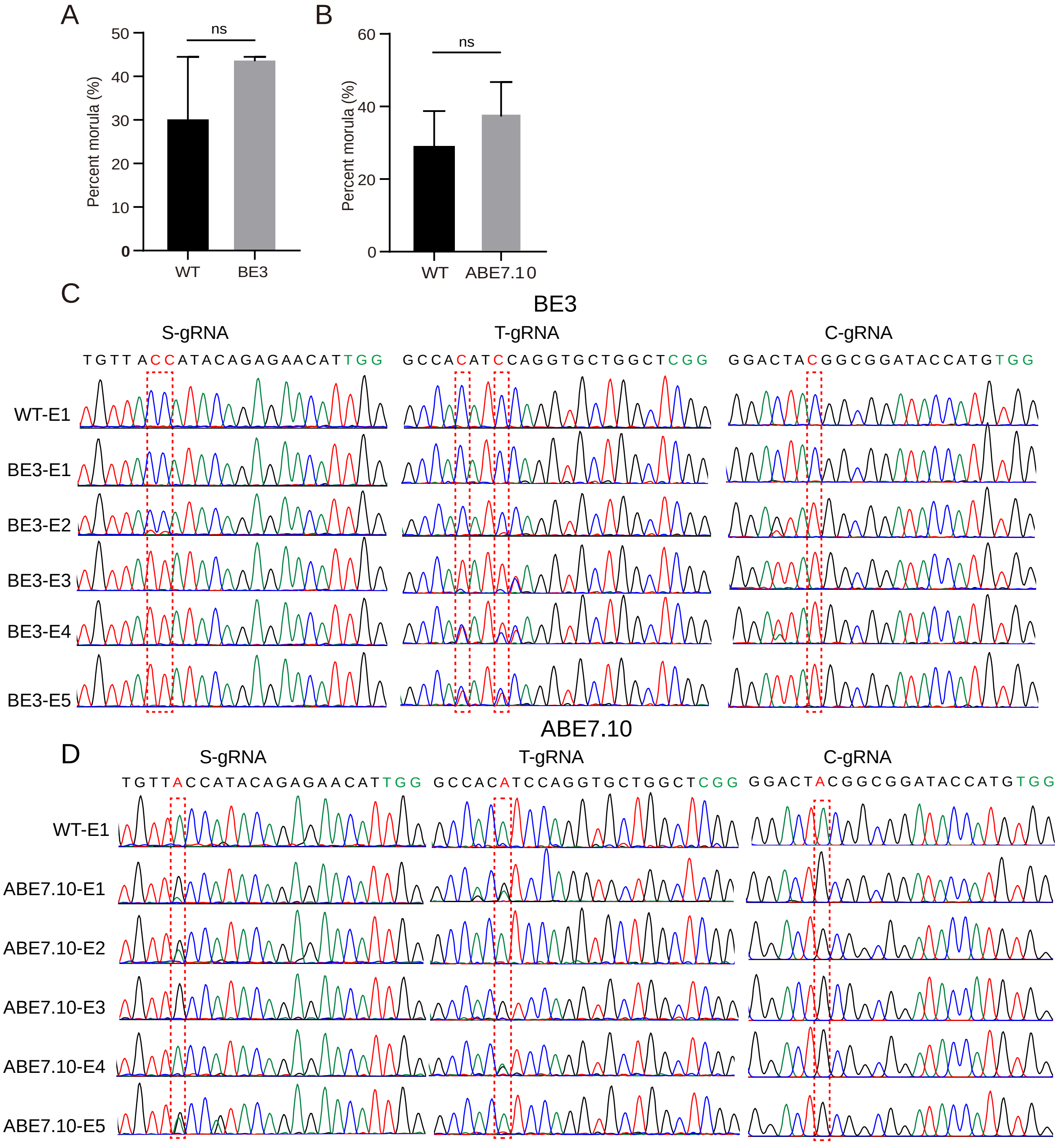
A
B
ns
Percent morula (%)
50
40
30
20
10
0
WT
BE3
60
40
20
0
ABE7.10
C
BE3
S-gRNA
T-gRNA
C-gRNA
TGTTACCATACAGAGAACATTGG
GCCACATCCAGGTGCTGGCTCGG
GGACTACGGCGGATACCATGTGG
WT-E1
BE3-E1
BE3-E2
BE3-E3
BE3-E4
BE3-E5
ABE7.10
D
TGTTACCATACAGAGAACATTGG
GCCACATCCAGGTGCTGGCTCGG
GGACTACGGCGGATACCATGTGG
ABE7.10-E1
ABE7.10-E2
ABE7.10-E3
ABE7.10-E4
ABE7.10-E5

**Figure S2. Application of multi-gene base editing using ABE7.10 and BE3**

A and B: Embryonic developmental rate to reach morula stage in the WT and ABE7.10 (A) or BE3 (B) group (Two replicates of 13-18 embryos per group). C. Representative Sanger sequencing results of ABE7.10-meidated base editing. D. Representative Sanger sequencing results of BE3-mediated base editing. The red letters and frames represent the edited sites. The green letters represent PAM sequence. S-gRNA: *SMAD4* sgRNA; T-gRNA: *TEAD4* sgRNA; C-gRNA: *CDX2* sgRNA.

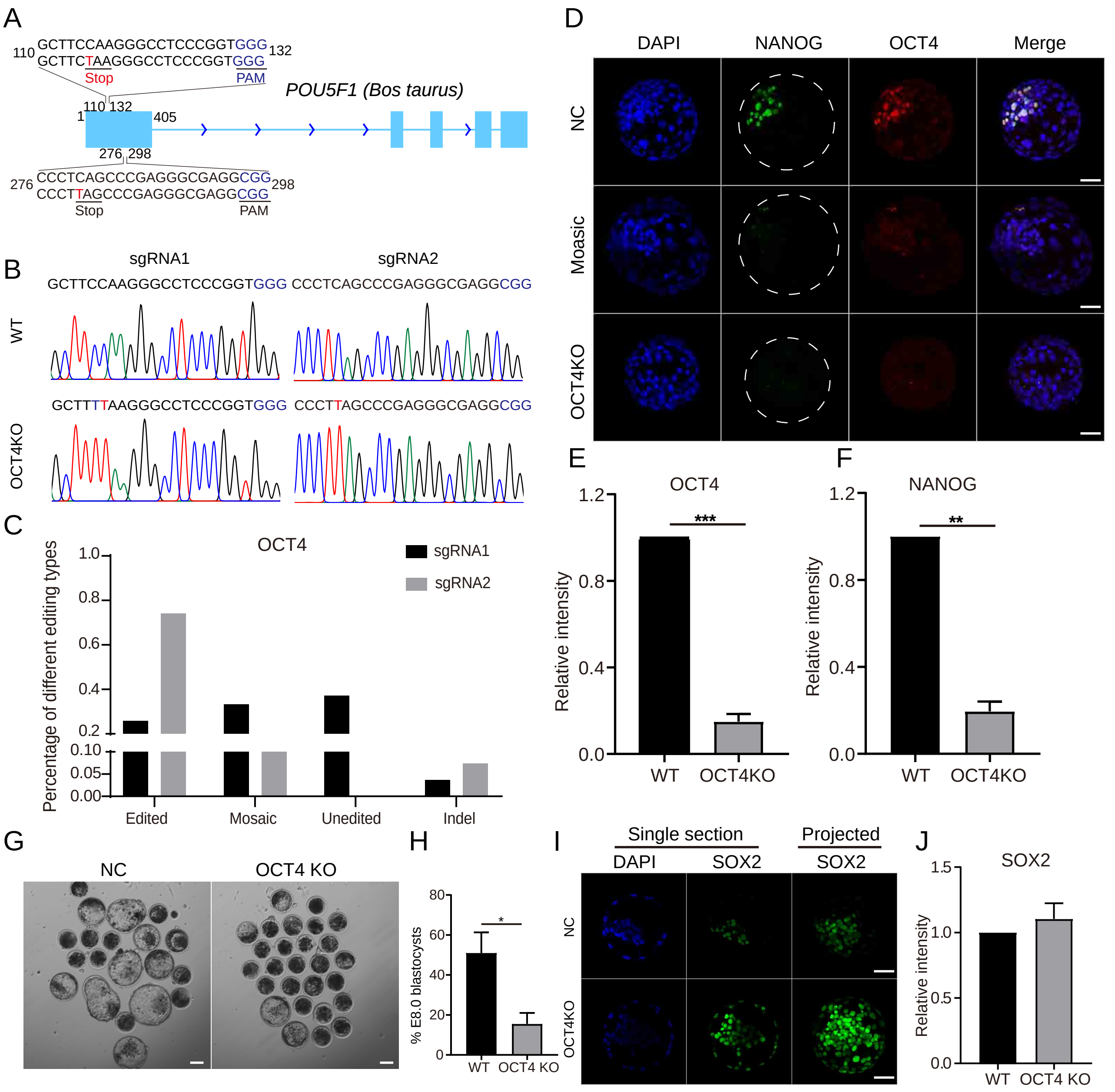

A
110 GCTTCCAAGGGCCTCCCGGTGGG 132
GCTTCTAAGGGCCTCCCGGTGGG
Stop
PAM
POU5F1 (Bos taurus)
1
110 132
405
276 298
276 CCCTCAGCCCGAGGGCGAGGCGG 298
CCCTTAGCCCGAGGGCGAGGCGG
Stop
PAM
B
sgRNA1
sgRNA2
GCTTCCAAGGGCCTCCCGGTGGG CCCTCAGCCCGAGGGCGAGGCGG
WT
GCTTTTAAGGGCCTCCCGGTGGG CCCTTAGCCCGAGGGCGAGGCGG
OCT4KO
C
OCT4
sgRNA1
sgRNA2
Percentage of different editing types
1.0
0.8
0.6
0.4
0.2
0.10
0.05
0.00
Edited
Mosaic
Unedited
Indel
D
DAPI
NANOG
OCT4
Merge
NC
Moasic
OCT4KO
E
OCT4
Relative intensity
1.2
0.8
0.4
0.0
***
WT
OCT4KO
F
NANOG
Relative intensity
1.2
0.8
0.4
0.0
**
WT
OCT4KO
G
NC
OCT4 KO
H
% E8.0 blastocysts
80
60
40
20
0
*
WT
OCT4 KO
I
Single section
Projected
DAPI
SOX2
SOX2
NC
OCT4KO
J
SOX2
Relative intensity
1.5
1.0
0.5
0.0
WT
OCT4 KO

**Figure S3. Effects of OCT4 knockout on bovine early embryonic development.**
A. Two sgRNAs designed to target *OCT4*. The red letters represent potential editing sites. B. Representative Sanger sequencing results. The red letters represent edited sites. C. Editing types of *OCT4* sgRNA1 and sgRNA2 (27 embryos were analyzed). D. Immunostaining detection of OCT4 and NANOG in WT and OCT4 KO groups (Two replicates of 4-6 blastocysts per group). Green: NANOG; Red: OCT4. Scale bar = 50 μm. E: OCT4 KO results in the decrease of blastocyst rate (Three replicates of 20-25 embryos per group). Scale bar = 100 μm. F. Immunostaining analysis of SOX2 expression and distribution (Two replicates of 4-6 blastocysts per group). Green: SOX2; Red: OCT4. Scale bar = 50 μm.

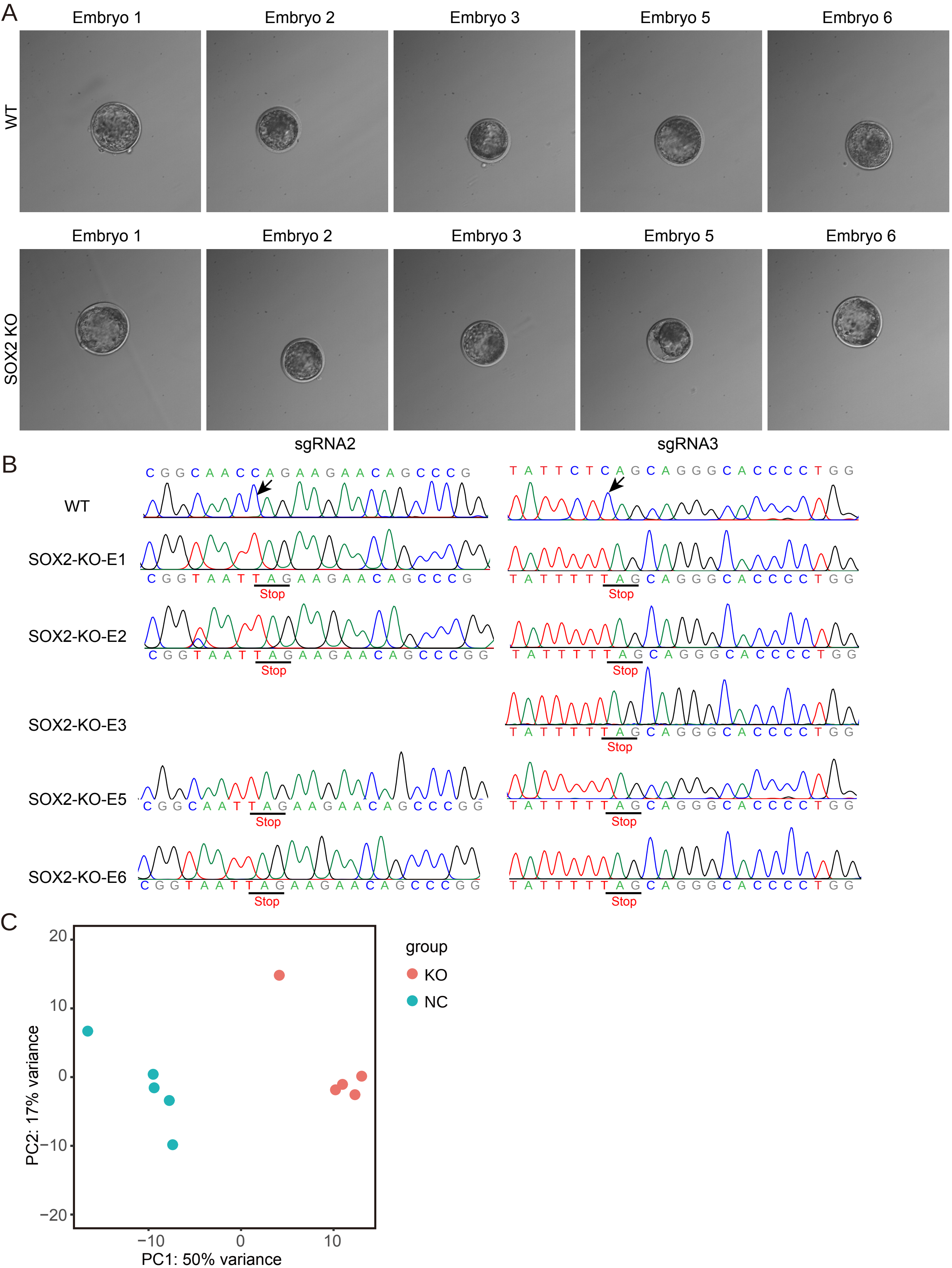
A
Embryo 1
Embryo 2
Embryo 3
Embryo 5
Embryo 6
WT
SOX2 KO
B
sgRNA2
sgRNA3
WT
SOX2-KO-E1
SOX2-KO-E2
SOX2-KO-E3
SOX2-KO-E5
SOX2-KO-E6
CGGCAACCAGAAGAACAGCCCG
TATTCTCAGCAGGGCACCCCTGG
CGGTAATTAGAAGAACAGCCCG
TATTTTTAGCAGGGCACCCCTGG
CGGTAATTAGAAGAACAGCCCGG
CGGCAATTAGAAGAACAGCCCGG
Stop
C
group
KO
NC
PC1: 50% variance
PC2: 17% variance

**Figure S4. Single blastocyst RNA sequencing**

A. Single blastocyst was collected (n=5 per group) to perform RNA sequencing. B. Validation of the genotypes of embryos used for RNA-sequencing in A. C. Principal component analysis (PCA) shows high correlation among samples in the same group.

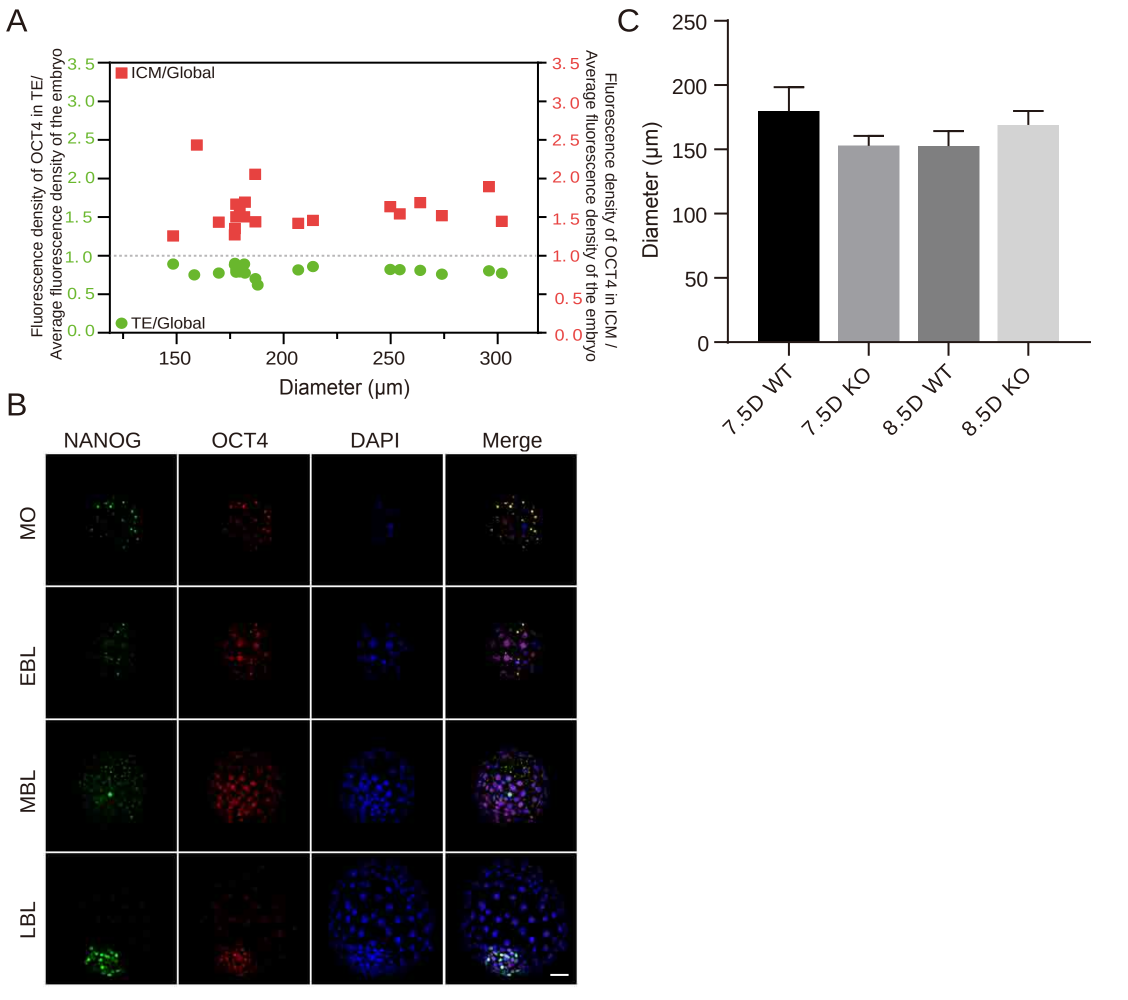
A
ICM/Global
TE/Global
Fluorescence density of OCT4 in TE/
Average fluorescence density of the embryo
Fluorescence density of OCT4 in ICM /
Average fluorescence density of the embryo
3.5
3.0
2.5
2.0
1.5
1.0
0.5
0.0
150
200
250
300
Diameter (μm)
B
NANOG
OCT4
DAPI
Merge
MO
EBL
MBL
LBL
C
Diameter (μm)
250
200
150
100
50
0
7.5D WT
7.5D KO
8.5D WT
8.5D KO

**Figure S5. SOX2 knockout did not affect blastocyst expansion**

A. Quantification analysis of the changes of OCT4 levels in TE and ICM as blastocyst expansion. TE: $SOX2^-$ cells; ICM: $SOX2^+$ cells. B. The dynamics of NANOG and OCT4 expression accompanied by the blastocyst expansion. Green: NANOG; Red: OCT4 (Two replicates of 3-5 blastocysts at different stages). Scale bar = 50 µm C. SOX2 knockout did not affect the blastocyst expansion at E7.5 or E8.5 (Two independent replicates of 12-17 embryos per group).

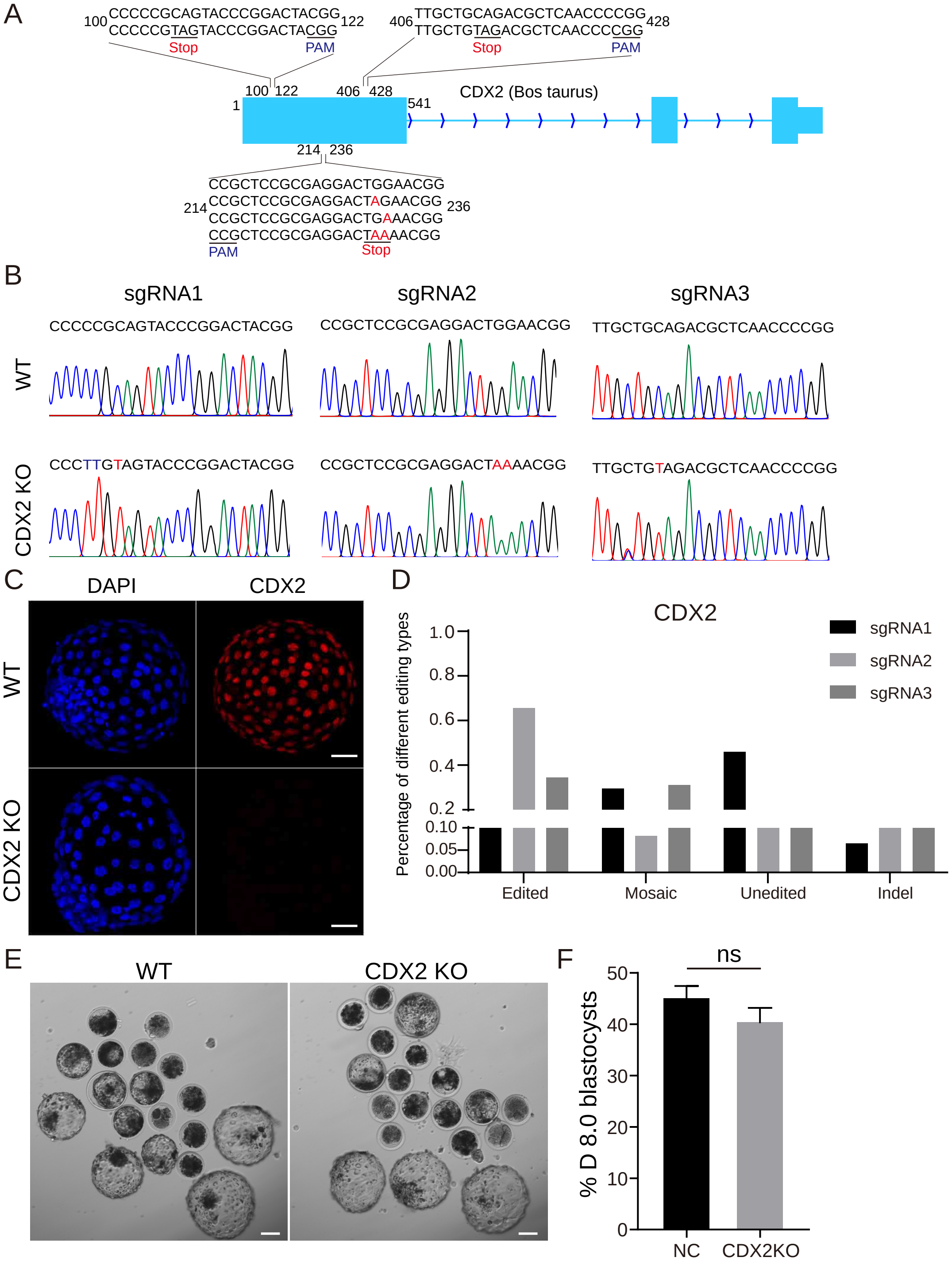
A
100 CCCCCGCAGTACCCGGACTACGG 122
CCCCCGTAGTACCCGGACTACGG
Stop
PAM
406 TTGCTGCAGACGCTCAACCCCGG 428
TTGCTGTAGACGCTCAACCCCGG
Stop
PAM
CDX2 (Bos taurus)
1
100 122
406 428
541
214 236
CCGCTCCGCGAGGACTGGAACGG
CCGCTCCGCGAGGACTAGAACGG
CCGCTCCGCGAGGACTGAAACGG
CCGCTCCGCGAGGACTAAAACGG
PAM
Stop
B
sgRNA1
sgRNA2
sgRNA3
WT
CCCCCGCAGTACCCGGACTACGG
CCGCTCCGCGAGGACTGGAACGG
TTGCTGCAGACGCTCAACCCCGG
CDX2 KO
CCCTTGTAGTACCCGGACTACGG
CCGCTCCGCGAGGACTAAAACGG
TTGCTGTAGACGCTCAACCCCGG
C
DAPI
CDX2
WT
CDX2 KO
D
CDX2
sgRNA1
sgRNA2
sgRNA3
Percentage of different editing types
1.0
0.8
0.6
0.4
0.2
0.10
0.05
0.00
Edited
Mosaic
Unedited
Indel
E
WT
CDX2 KO
F
ns
% D 8.0 blastocysts
50
40
30
20
10
0
NC
CDX2KO

**Figure S6. CDX2 KO did not affect bovine early embryonic development.**

A. sgRNAs designed to target CDX2. B. Representative Sanger sequencing results of CDX2 editing. The red letters represent edited sites. C. Immunostaining detection of CDX2 (Three replicates of 5-8 blastocysts per group). Red: CDX2. Scale bar = 50 μm. D. Editing types analysis using *CDX2* sgRNA1, sgRNA2 and sgRNA3 (61 embryos were detected). E and F. CDX2 KO has no effect on the rate of blastocyst formation (Ten replicates of 20-25 embryos per group). Scale bar = 100 μm.

**Table S1. The synthesis of sgRNAs sequence**

| Gene name | Gene ID | sgRNA | Exon | Sequence（5' – 3'） |
|---|---|---|---|---|
| *SMAD4* | 540248 | S-gRNA | 1 | FP：CACCTGTTACCATACAGAGAACAT<br>RP：AAACATGTTCTCTGTATGGTAACA |
| *TEAD4* | 526771 | T-gRNA | 5 | FP：CACCGCCACATCCAGGTGCTGGCT<br>RP：AAACAGCCAGCACCTGGATGTGGC |
| *CDX2* | 618679 | C-gRNA | 1 | FP：CACCGGACTACGGCGGATACCATG<br>RP：AAACCATGGTATCCGCCGTAGTCC |
| *CDX2* | 618679 | sgRNA1 | 1 | FP：CACCCCCCCGCAGTACCCGGACTA<br>RP：AAACTAGTCCGGGTACTGCGGGGG |
| *CDX2* | 618679 | sgRNA2 | 1 | FP：CACCCCGTTCCAGTCCTCGCGGAG<br>RP：AAACCTCCGCGAGGACTGGAACGG |
| *CDX2* | 618679 | sgRNA3 | 1 | FP：CACCTTGCTGCAGACGCTCAACCC<br>RP：AAACGGGTTAGACGTCTGCAGCAA |
| *OCT4* | 282316 | sgRNA1 | 1 | FP：CACCGCTTCCAAGGGCCTCCCGGT<br>RP：AAACACCGGGAGGCCCTTGGAAGC |
| *OCT4* | 282316 | sgRNA2 | 1 | FP：CACCCCCTCAGCCCGAGGGCGAGG<br>RP：AAACCCTCGCCCTCGGGCTGAGGG |
| *SOX2* | 784383 | sgRNA1 | 1 | FP：CACCCCGCAGCAAACTTCGGGGGG<br>RP：AAACCCCCCCGAAGTTTGCTGCGG |
| *SOX2* | 784383 | sgRNA2 | 1 | FP：CACCCGGCAACCAGAAGAACAGCC<br>RP：AAACGGCTGTTCTTCTGGTTGCCG |
| *SOX2* | 784383 | sgRNA3 | 1 | FP：CACCTATTCTCAGCAGGGCACCCC<br>RP：AAACGGGGTGCCCTGCTGAGAATA |

**Table S2. The primers information of sgRNA template for in vitro transcription**

| sgRNA | Primers' name | Sequence（5' – 3'） |
|---|---|---|
| S-gRNA | S-g1-T7-F | TTAATACGACTCACTATAGTGTTACCATACAGAGAACAT |
| T-gRNA | T-g1-T7-F | TTAATACGACTCACTATAGCCACATCCAGGTGCTGGCT |
| C-gRNA | C-g1-T7-F | TTAATACGACTCACTATAGGACTACGGCGGATACCATG |
| CDX2-sgRNA1 | C-g2-T7-F | TTAATACGACTCACTATAGCCCCCGCAGTACCCGGACTA |
| CDX2-sgRNA2 | C-g3-T7-F | TTAATACGACTCACTATAGCCGTTCCAGTCCTCGCGGAG |
| CDX2-sgRNA3 | C-g4-T7-F | TTAATACGACTCACTATAGTTGCTGCAGACGCTCAACCC |
| OCT4-sgRNA1 | OCT4-g1-T7-F | TAATACGACTCACTATAGGCTTCCAAGGGCCTCCCGGT |

| sgRNA | Primers' name | Sequence（5' – 3'） |
|---|---|---|
| OCT4-sgRNA2 | OCT4-g2-T7-F | TAATACGACTCACTATAGGCCCTCAGCCCGAGGGCGAGG |
| SOX2-sgRNA1 | SOX2-g1-T7-F | TAATACGACTCACTATAGGCCGCAGCAAACTTCGGGGGG |
| SOX2-sgRNA2 | SOX2-g2-T7-F | TAATACGACTCACTATAGGCGGCAACCAGAAGAACAGCC |
| SOX2-sgRNA3 | SOX2-g3-T7-F | TAATACGACTCACTATAGGTATTCTCAGCAGGGCACCCC |
| | sgRNA-R | AAAAGCACCGACTCGGTGCC |

**Table S3. Nested PCR primer sequences for preparing Sanger sequencing samples**

| Gene | Gene ID | Target sgRNAs | Primers' name | Sequence（5' – 3'） |
| --- | --- | --- | --- | --- |
| *SMAD4* | 540248 | S-gRNA | S-g1-1 | FP：TCTAACAATTTTCCTTGCAAC<br>RP：CCTGTATTGATAATATCTGTGCT |
| | | | S-g1-2 | FP：TCCGAAAGATCAAAATTGCT<br>RP：TACAGTATCTAAAGAGACGGAG |
| *TEAD4* | 526771 | T-gRNA | T-g1-1 | FP：CTGAGAGGGTGCTGTGTCTC<br>RP：AACGTCCAGTCCCAGAGAGT |
| | | | T-g1-2 | FP：GGACCGCACTTCTGTTTAGC<br>RP：TGGGGCTTACAGGGTTACAG |
| *CDX2* | 618679 | C-gRNA<br>CDX2-sgRNA1<br>CDX2-sgRNA2<br>CDX2-sgRNA3 | C-g-1 | FP：ATGGTGAGGTTCGCCGTC<br>RP：GCTTTACACTGAACGCGGCT |
| | | | C-g-2 | FP：TACGTGAGCTACCTCCTGGAC<br>RP：CCCCGCTATCCCCTACTCA |
| *OCT4* | 282316 | OCT4-sgRNA1<br>OCT4-sgRNA2 | OCT4-g-1 | FP：GCCAGAGGTCAAGGCTAGTG<br>RP：CTCACCTGCGGTTCTCTCTT |
| | | | OCT4-g-2 | FP：CGGGACACCTCGCTTCTGAC<br>RP：CTTCGCCTGCTCCCTTCCTG |
| *SOX2* | 784383 | SOX2-sgRNA1<br>SOX2-sgRNA2<br>SOX2-sgRNA3 | SOX2-g-1 | FP：CCCCCAAATTATTCTTCGCCTG<br>RP：CTTTGAAAATGTCTCCCCCG |
| | | | SOX2-g-2 | FP：GCCCGCATGTACAACATGATGG<br>RP：GGGCAGTGTGCCGTTAATGG |

**Table S4. The predicted potential 6 off-target sites for *SMAD4***

| Name | Chromosome | Sequence information |
| --- | --- | --- |
| OFF 1 | 2 | TGTTACCATAGCAGAGGACAT |
| OFF 2 | 23 | TTTTACCATAAAGAGAACTT |
| OFF 3 | 11 | TCTTACCATTCAGAGAAGCAT |
| OFF 4 | 7 | TGTTTGCATACAGAGAAGAT |

| | | |
|---|---|---|
| OFF 5 | 9 | TGATCCATACAGAGAAGAT |
| OFF 6 | 1 | TGTTACCAGCCAGAGAACACT |

**Table S5 PCR primer sequences for preparing targeted next generation sequencing samples**

| Amplified sites | Chromosome | Primers' name | Sequence（5' – 3'） |
|---|---|---|---|
| S-gRNA-ON | 24 | S-g1-ON-F1 | ATCACGGGAGAGAGTGAAACATTCGCC |
| | | S-g1-ON-F2 | TTAGGCGGAGAGAGTGAAACATTCGCC |
| | | S-g1-ON-F3 | ACAGTGGGAGAGAGTGAAACATTCGCC |
| | | S-g1-ON-F4 | CAGATCGGAGAGAGTGAAACATTCGCC |
| | | S-g1-ON-F5 | TAGCTTGGAGAGAGTGAAACATTCGCC |
| | | S-g1-ON-F6 | GGCTACGGAGAGAGTGAAACATTCGCC |
| | | S-g1-ON-R1 | CGATGTAAAGAGACGGAGCAAGCATAAA |
| | | S-g1-ON-R2 | TGACCAAAAGAGACGGAGCAAGCATAAA |
| | | S-g1-ON-R3 | GCCAATAAAGAGACGGAGCAAGCATAAA |
| | | S-g1-ON-R4 | ACTTGAAAAGAGACGGAGCAAGCATAAA |
| | | S-g1-ON-R5 | GATCAGAAAGAGACGGAGCAAGCATAAA |
| S-gRNA-OFF1 | 2 | S-g1-OFF1-F1 | ATCACGTCTTTGTGCAGAATGGGGGT |
| | | S-g1-OFF1-F2 | TTAGGCTCTTTGTGCAGAATGGGGGT |
| | | S-g1-OFF1-F3 | ACAGTGTCTTTGTGCAGAATGGGGGT |
| | | S-g1-OFF1-F4 | CAGATCTCTTTGTGCAGAATGGGGGT |
| | | S-g1-OFF1-F5 | TAGCTTTCTTTGTGCAGAATGGGGGT |
| | | S-g1-OFF1-F6 | GGCTACTCTTTGTGCAGAATGGGGGT |
| | | S-g1-OFF1-R1 | CGATGTGATGATTCCCGAGTGGGAGC |
| | | S-g1-OFF1-R2 | TGACCAGATGATTCCCGAGTGGGAGC |
| | | S-g1-OFF1-R3 | GCCAATGATGATTCCCGAGTGGGAGC |
| | | S-g1-OFF1-R4 | ACTTGAGATGATTCCCGAGTGGGAGC |
| | | S-g1-OFF1-R5 | GATCAGGATGATTCCCGAGTGGGAGC |
| S-gRNA-OFF2 | 23 | S-g1-OFF2-F1 | ATCACGAATTCACATCATGCTTTTGTTTCAT |
| | | S-g1-OFF2-F2 | TTAGGCAATTCACATCATGCTTTTGTTTCAT |
| | | S-g1-OFF2-F3 | ACAGTGAATTCACATCATGCTTTTGTTTCAT |
| | | S-g1-OFF2-F4 | CAGATCAATTCACATCATGCTTTTGTTTCAT |

| Amplified sites | Chromosome | Primers' name | Sequence（5' – 3'） |
|---|---|---|---|
| | | S-g1-OFF2-F5 | TAGCTTAATTCACATCATGCTTTTGTTTCAT |
| | | S-g1-OFF2-F6 | GGCTACAATTCACATCATGCTTTTGTTTCAT |
| | | S-g1-OFF2-R1 | CGATGTGCACATGCAGAGATATTTTCTAAGT |
| | | S-g1-OFF2-R2 | TGACCAGCACATGCAGAGATATTTTCTAAGT |
| | | S-g1-OFF2-R3 | GCCAATGCACATGCAGAGATATTTTCTAAGT |
| | | S-g1-OFF2-R4 | ACTTGAGCACATGCAGAGATATTTTCTAAGT |
| | | S-g1-OFF2-R5 | GATCAGGCACATGCAGAGATATTTTCTAAGT |
| S-gRNA-OFF3 | 11 | S-g1-OFF3-F1 | ATCACGGGCGCCTTGATTAATACCCC |
| | | S-g1-OFF3-F2 | TTAGGCGGCGCCTTGATTAATACCCC |
| | | S-g1-OFF3-F3 | ACAGTGGGCGCCTTGATTAATACCCC |
| | | S-g1-OFF3-F4 | CAGATCGGCGCCTTGATTAATACCCC |
| | | S-g1-OFF3-F5 | TAGCTTGGCGCCTTGATTAATACCCC |
| | | S-g1-OFF3-F6 | GGCTACGGCGCCTTGATTAATACCCC |
| | | S-g1-OFF3-R1 | CGATGTGCAACTACCGTGAGAGTTCA |
| | | S-g1-OFF3-R2 | TGACCAGCAACTACCGTGAGAGTTCA |
| | | S-g1-OFF3-R3 | GCCAATGCAACTACCGTGAGAGTTCA |
| | | S-g1-OFF3-R4 | ACTTGAGCAACTACCGTGAGAGTTCA |
| | | S-g1-OFF3-R5 | GATCAGGCAACTACCGTGAGAGTTCA |
| S-gRNA-OFF4 | 7 | S-g1-OFF4-F1 | ATCACGACCAGACGAGGTACTGTAACTTT |
| | | S-g1-OFF4-F2 | TTAGGCACCAGACGAGGTACTGTAACTTT |
| | | S-g1-OFF4-F3 | ACAGTGACCAGACGAGGTACTGTAACTTT |
| | | S-g1-OFF4-F4 | CAGATCACCAGACGAGGTACTGTAACTTT |
| | | S-g1-OFF4-F5 | TAGCTTACCAGACGAGGTACTGTAACTTT |
| | | S-g1-OFF4-F6 | GGCTACACCAGACGAGGTACTGTAACTTT |
| | | S-g1-OFF4-R1 | CGATGTCCGGAGGAGCAGACTAGTGAT |
| | | S-g1-OFF4-R2 | TGACCACCGGAGGAGCAGACTAGTGAT |
| | | S-g1-OFF4-R3 | GCCAATCCGGAGGAGCAGACTAGTGAT |
| | | S-g1-OFF4-R4 | ACTTGACCGGAGGAGCAGACTAGTGAT |
| | | S-g1-OFF4-R5 | GATCAGCCGGAGGAGCAGACTAGTGAT |
| S-gRNA-OFF5 | 9 | S-g1-OFF5-F1 | ATCACGTGAGGAGAATGCCAACTTGTTC |
| | | S-g1-OFF5-F2 | TTAGGCTGAGGAGAATGCCAACTTGTTC |

| Amplified sites | Chromosome | Primers' name | Sequence（5' – 3'） |
|---|---|---|---|
| | | S-g1-OFF5-F3 | ACAGTGTGAGGAGAATGCCAACTTGTTC |
| | | S-g1-OFF5-F4 | CAGATCTGAGGAGAATGCCAACTTGTTC |
| | | S-g1-OFF5-F5 | TAGCTTTGAGGAGAATGCCAACTTGTTC |
| | | S-g1-OFF5-F6 | GGCTACTGAGGAGAATGCCAACTTGTTC |
| | | S-g1-OFF5-R1 | CGATGTCTGTATTTCAAAATGTAGGACCGT |
| | | S-g1-OFF5-R2 | TGACCACTGTATTTCAAAATGTAGGACCGT |
| | | S-g1-OFF5-R3 | GCCAATCTGTATTTCAAAATGTAGGACCGT |
| | | S-g1-OFF5-R4 | ACTTGACTGTATTTCAAAATGTAGGACCGT |
| | | S-g1-OFF5-R5 | GATCAGCTGTATTTCAAAATGTAGGACCGT |
| S-gRNA-OFF6 | 1 | S-g1-OFF6-F1 | ATCACGCACGGTCCAGTAGGTTCCCA |
| | | S-g1-OFF6-F2 | TTAGGCCACGGTCCAGTAGGTTCCCA |
| | | S-g1-OFF6-F3 | ACAGTGCACGGTCCAGTAGGTTCCCA |
| | | S-g1-OFF6-F4 | CAGATCCACGGTCCAGTAGGTTCCCA |
| | | S-g1-OFF6-F5 | TAGCTTCACGGTCCAGTAGGTTCCCA |
| | | S-g1-OFF6-F6 | GGCTACCACGGTCCAGTAGGTTCCCA |
| | | S-g1-OFF6-R1 | CGATGTCCTTCCGATTCCACGCAGT |
| | | S-g1-OFF6-R2 | TGACCACCTTCCGATTCCACGCAGT |
| | | S-g1-OFF6-R3 | GCCAATCCTTCCGATTCCACGCAGT |
| | | S-g1-OFF6-R4 | ACTTGACCTTCCGATTCCACGCAGT |
| | | S-g1-OFF6-R5 | GATCAGCCTTCCGATTCCACGCAGT |

### Table S6：Antibody information

| Name | Host | Company | Catalog Number | Application |
|---|---|---|---|---|
| SOX2 | Rat | eBioscience™ | 14-9811-82 | IF (1:200) |
| CDX2 | Mouse | BioGenex | CDX2-88 | IF (1:20) |
| GATA3 | Rabbit | Abcam | ab199428 | IF (1:200) |
| OCT4 | Rabbit | Abcam | Ab181557 | IF (1:200) |
| NANOG | Mouse | eBioscience™ | 14-5768-82 | IF (1:100) |
| SOX17 | Goat | Bio-techne | AF1924 | IF (1:200) |
